# Electrical Spinal Imaging (ESI): Analysing spinal cord activity with non-invasive, high-resolution mapping

**DOI:** 10.1101/2025.03.17.643665

**Authors:** G Gabrieli, R Somervail, A Mouraux, M Leandri, P Haggard, GD Iannetti

**Affiliations:** Neuroscience and Behaviour Laboratory, Italian Institute of Technology, Rome, Italy; Institute of Neuroscience, Université Catholique de Louvain, Brussels, Belgium; DINOGMI, Laboratory of Clinical and Experimental Electrophysiology. University of Genova, Genova, Italy; Department of Neuroscience Physiology and Pharmacology, University College London, London, UK

## Abstract

The spinal cord is the key bridge between the brain and the body. However, scientific understanding of healthy spinal cord function has historically been limited because noninvasive measures of its neural activity have proven exceptionally challenging.

In this work, we describe a novel recording and analysis approach to obtain non-invasive, high-resolution images of the electrical activity of the spinal cord in humans (Electrical Spinal Imaging, ESI). ESI is analytically simple, easy to implement, and data-driven: it does not involve template-based strategies prone to produce spurious signals. Using this approach we provide a detailed description and physiological characterization of the spatiotemporal dynamics of the peripheral, spinal and cortical activity elicited by somatosensory stimulation. We also demonstrate that attention modulates post-synaptic activity at spinal cord level.

Our method has enabled four new insights regarding spinal cord activity. (1) We identified three distinct responses in the time domain: sP9, sN13 and sP22. (2) The sP9 is a traveling wave reflecting the afferent volley entering the spinal cord through the dorsal root. (3) In contrast, the sN13 and sP22 reflect segmental post-synaptic activity. (4) While the sP9 response is first seen on the dorsal electrodes ipsilateral to the stimulated side, the sN13 and sP22 were not lateralised with respect to the side of stimulation. (5) Unimodal attention strongly modulates the amplitude of the sP22, but not that of the sP9 and sN13 components.

The proposed method offers critical insights into the spatiotemporal dynamics of somatosensory processing within the spinal cord, paving the way for precise non-invasive functional monitoring of the spinal cord in basic and clinical neurophysiology.

## Introduction

Electrical brain potentials elicited by somatosensory stimulation are widely employed in basic research and clinical practice. They are routinely used to verify the functional integrity of afferent pathways (Carter & Stevens, 2009), and have allowed obtaining a wealth of information about the spatiotemporal dynamics of the cortical processing of somatosensory input (Mountcastle, 2005).

A similarly reliable, direct and non-invasive readout of neural activity in the spinal cord *in vivo* would be equally informative, and has therefore been considered a key aim in sensory physiology (Eisen, 1986). Recent advances in fMRI technology have allowed recording blood-oxygen level dependent (BOLD) signals arising in the spinal cord. However, in addition to the severe technical challenges posed by cardiac and respiration induced movement artifacts, differences in magnetic susceptibilities of nearby tissues, cerebrospinal fluid inflow artifacts (Eippert et al., 2017; Giove et al., 2004; Maieron et al., 2007), the most important pitfall is the fact that BOLD signal integrates neural activities over several seconds, thus intrinsically limiting the ability to resolve neural events in time (Logothetis, 2003). Thus, despite allowing localization of spinal responses during motor or somatosensory tasks (Eippert et al., 2017; Maieron et al., 2007; Stroman, 2005) there has been a continuous effort to combine fMRI-based investigations with a more direct readout of the spinal electrical activity.

Recording the electrical activity of the spinal cord non-invasively is, however, not an easy task either. The first issue is neurophysiological: it is unclear whether the degree of regular arrangement of spinal neurons allows a reliable summation of post-synaptic potentials (Willis & Coggeshall, 2004). Indeed, while the scalp EEG results from the summation of synchronous activity from thousands of spatially-aligned cortical neurons (da Silva, 2022), neurons within the cord are neither as numerous or as aligned (Willis & Coggeshall, 2004). Second, surface recordings from the neck or dorsum skin are severely affected by muscular artifacts orders of magnitude larger than any neural signal. Third, cardiac activity similarly generates a massive artifact that, however, has the advantage of being predictable. Finally, the electrical activity of the somatosensory volleys ascending in the dorsal roots and intrinsic spinal tracts overlaps in space and time with the post-synaptic activity of the cord. All these factors result in an extremely low signal-to-noise ratio, unless epidural or intraparenchymal recordings are performed (Ertekin, 1978).

For this reason, recent attempts at recording the electrospinal activity have resorted to the use of selective spatiotemporal filtering and template-matching strategies to enhance the signal-to-noise ratio, such as denoising separation of sources (DSS; Chander et al., 2022) or canonical correlation analysis (CCA; Nierula et al., 2024). These approaches, however, are biased and prone to the risk of creating spurious responses (Hu & Iannetti, 2016; Li et al., 2009; Mayhew et al., 2006; Woody, 1967).

In this work we propose a novel methodology (Electrical Spinal Imaging, ESI) that couples (1) high-density (63-channel) recording of the electrospinal activity with (2) a simple analysis pipeline based on conservative signal cleaning that minimises the alterations of the spatiotemporal properties of the signal and does not rely on operator-based decisions (Figure 1).

**Figure 1.**
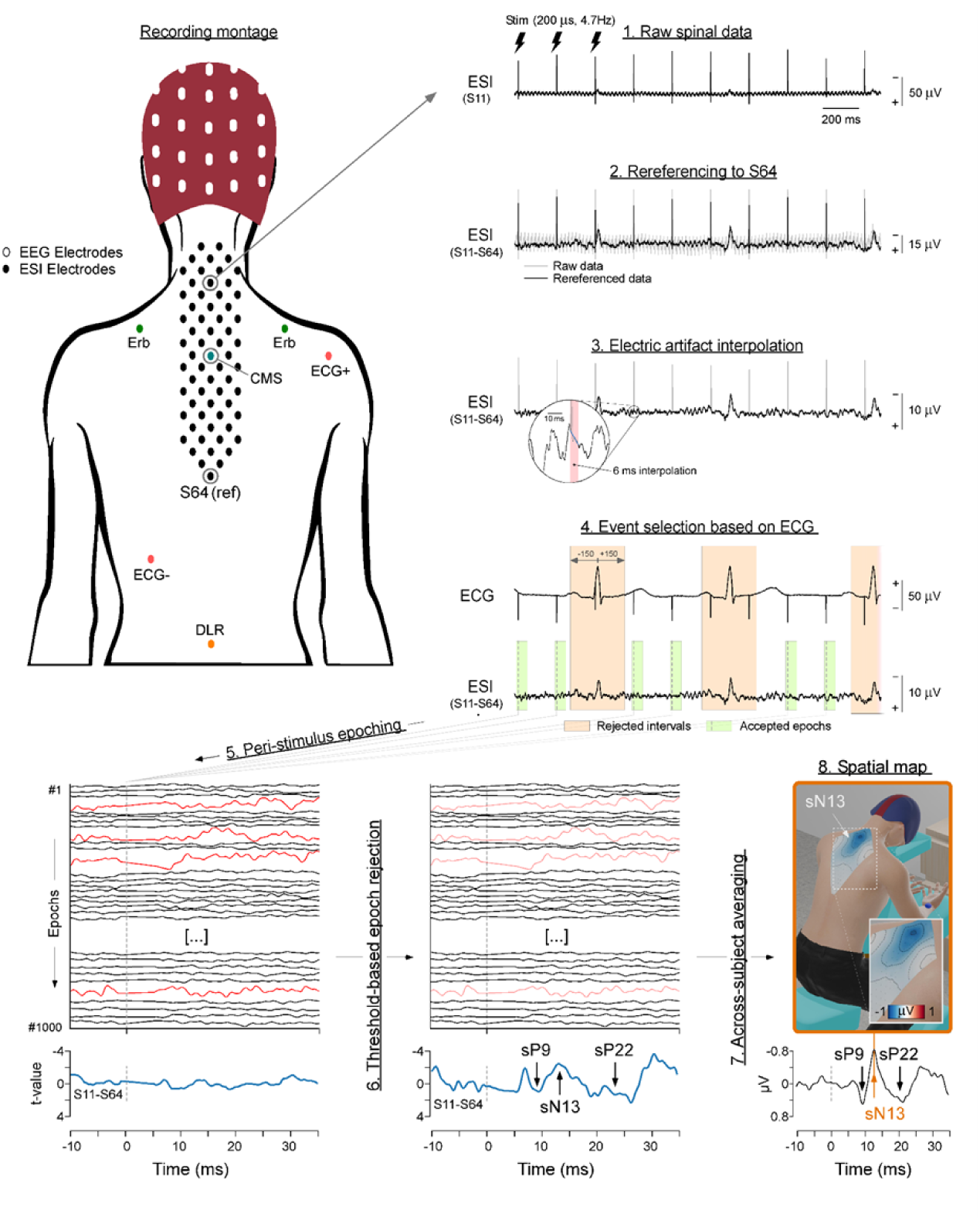
Recording setup and data analysis. *Top left panel:* Schematic of the position of the electrodes to record EEG (white), ESI (black), ECG (red), somatosensory activity at Erb’s points (green), and the positions of CMS (blue) and DRL (orange) electrodes. Note that Erb and ECG electrodes were placed on the chest, while ESI electrodes were placed on the back. *Top right and bottom panels:* Flowchart describing the analysis procedure. (1 and 2) Raw ESI signals are first re-referenced to the most caudal dorsal electrode (S64). (3) The artifact caused by the electrical stimulation of the median nerve is removed by linear interpolation. (4) The ECG is used to identify ESI time windows contaminated by the QRS complex (orange). This allows subsequent selection of ESI time windows to retain. (5) These time windows are epoched around the somatosensory stimulus. (6) An amplitude-based threshold is used to identify and remove artifactual epochs (red). (7) Resulting artifact-free epochs are averaged across stimuli of each block. Subject-level average waveforms are subsequently averaged across participants. (8) Images of the spatial distribution of spinal cord activity are calculated by spline interpolation across ESI electrodes. The figurine depicts spinal cord activity at the latency of the N13 wave.

**Figure 2.**
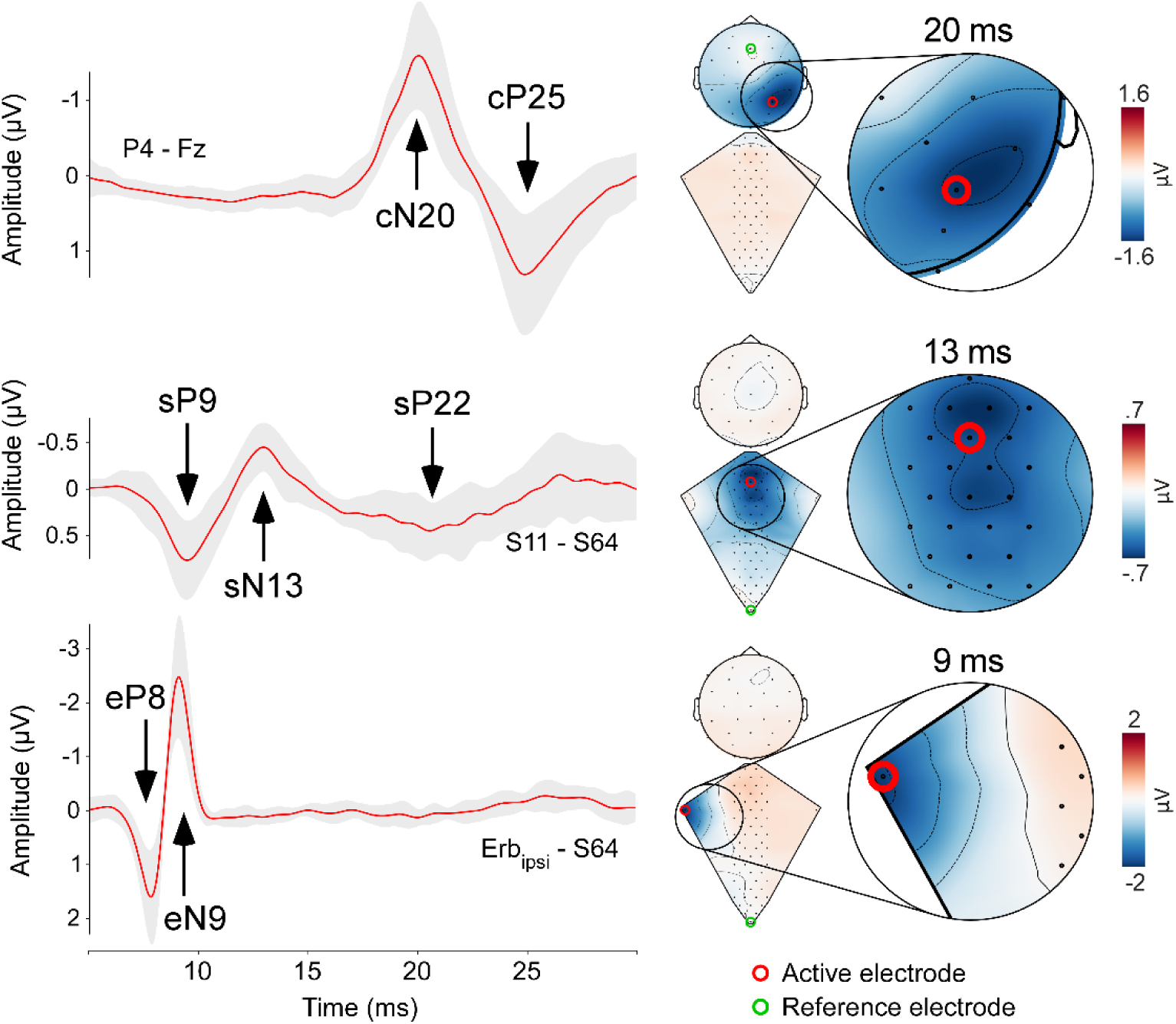
Neural activity along the somatosensory pathways in response to transcutaneous electrical stimulation of the median nerve at the wrist. Responses recorded from the ipsilateral brachial plexus (eP8 and eN9, bottom row), the spinal cord (sP9, sN13 and sP22, middle row), and the cerebral cortex (cN20-cP25, top row). Left column: red waveforms are group-level averages, while the gray shade represents the standard deviation (±1 standard deviation). Right columns: dorsum and scalp maps obtained using cubic interpolation of multielectrode recordings (dorsum: 63 electrodes; scalp: 32 electrodes). Maps are shown at the latency of the eN9, sN13, and cN20 peaks. Red circles show the active electrodes from which waveforms shown in the left column are extracted. Green circles show the reference electrode for both waveforms and maps: dorsal recordings are referenced to the most caudal electrode (S64), whereas scalp recordings are referenced to Fz. Enlargements show areas of maximal response amplitude.

Using this approach we provide a novel physiological characterization of the spatiotemporal dynamics of the peripheral, spinal and cortical activity elicited by somatosensory stimulation. We demonstrate a specific circuit mediating the attentional modulation of post-synaptic activity at spinal cord level.

## Methods

### Participants

16 healthy human participants (mean±SD age 28.9±5.1 years, range 19-37 years, 9 females) gave written consent before taking part in the study. Procedures were approved by the ethical committee of the Fondazione Santa Lucia, Rome, Italy.

### Experimental design

Experiments took place in a dim, quiet, temperature-controlled room. Participants were seated in a massage chair with their neck supported in a flexed position via a face cradle, and their arms kept semi-flexed and resting on two dedicated supports. Participants were asked to keep their gaze stable and minimize spontaneous eye blinks.

The experiment consisted of 8 successive blocks, each lasting approximately 4 minutes. Each block consisted of 1,000 somatosensory stimuli delivered at 4.75 Hz. The interval between consecutive blocks was approximately 2 minutes. In 4 blocks stimuli were delivered to the left median nerve. In the other 4 blocks stimuli were delivered to the right median nerve. The stimulated side was alternated each block. In half of the participants the first block entailed the stimulation of the right median nerve.

In each block, the stream of somatosensory stimuli was interrupted by a few omissions of individual stimuli. Omissions were randomly distributed across the block, subject to the constraint that at least 5 consecutive stimuli had to be delivered between omissions. The number of omissions ranged from 2 to 25 per block, with a rectangular distribution. With the standard interstimulus interval being 211 ms, the interval between two stimuli separated by an omission was 420 ms. Before each block, participants were instructed to either (1) count the number of omissions (*omission counting*), or (2) memorize a sequence of eight capital letters presented on an A4 page for a maximum of 2 minutes prior to starting the block (*letter memorization*). At the end of each block, participants were required to report either the number of omissions or the sequence of letters. There were 2 omissions counting and 2 letter memorization blocks per stimulation side. In half of the participants the first block entailed the omission counting task.

### Sensory stimuli

Somatosensory stimuli were constant-current, biphasic square-wave electrical pulses (200 μs duration; DS8R, Digitimer, UK) delivered using a surface bipolar electrode (0.5 cm diameter, 3 cm interelectrode distance, cathode sited proximally) placed over the median nerve of either the left or the right wrist. The intensity of somatosensory stimulation was initially adjusted for each participant to elicit a reproducible thumb muscle twitch, and then delivered 10% above this motor threshold during the stimulation blocks.

### Spinal recording

The electrical activity of the spinal cord was recorded using 63 active electrodes placed on the skin of the back (dorsal electrodes; Figure 1). Electrodes were arranged in 7 rostrocaudal columns: 1 column aligned on the spinal midline, and 6 parasagittal columns (3 to the right and 3 to the left side of the midline). The distance between adjacent columns was 7.5 mm. From left to right, the number of electrodes composing the seven columns was: 8, 9, 10, 9, 10, 9, 8. The distance between consecutive electrodes within each column was 3 cm. The two most rostral electrodes, which belonged to columns 3 and 5, were placed at the level of C2, each 7.5 mm from the midline (Figure 1). The remaining electrodes were progressively placed according to the above interelectrode distances. Thus, the entire array covered the skin spanning from the 2^nd^ cervical (C2) to approximately the 8^th^ thoracic (T8) vertebra. Two additional electrodes were placed on the left and right Erb’s point, to record the activity from the brachial plexus. The CMS (Common Mode Sense) and DRL (Driven Right Leg) electrodes were placed on the midline, one at the center of the array and the other at the level of the 3^rd^ lumbar vertebra (L3), respectively.

Signals were amplified, and digitized at a sampling rate of 4,096 Hz (Biosemi Active-2 system). The same amplifier was used to simultaneously record the EEG and the ECG (see section “EEG and ECG recordings”). Biosemi active electrode offsets were kept below 20 mV.

### EEG and ECG recordings

The EEG was recorded using 32 active electrodes placed on the scalp according to the International 10-20 system. The ECG was recorded using two surface electrodes placed on the anterior aspect of the right shoulder and at the level of the seventh intercostal space on the left mid clavicular line.

### Data processing

All electrophysiological data were preprocessed and analysed in Python (3.10.12) using the *mne* (Gramfort et al., 2013), *numpy* (Harris et al., 2020), and *scipy* (Virtanen et al., 2020) packages.

#### ECG processing

Continuous ECG signals were first band-pass filtered at 8-16Hz. Then, the latency of the R peak of each QRS complex was identified using a validated method implemented in mne (Gramfort et al., 2013). This procedure allowed removal of stimulation epochs from both electrocortical and electrospinal data that occurred within ±150 ms of the R peak, as described below.

#### ESI and EEG processing

A series of preprocessing steps was adopted to yield ESI and EEG epochs free from the large artifacts consequent to (1) electrical stimulation, (2) heartbeat, (3) muscular activity, and (4) other sources of noise. The overall ESI procedure is depicted in Figure 1.

Continuous ESI and EEG data were first re-referenced to the most caudal spinal electrode (S64; Figure 1), and to Fz, respectively. Then, electrical stimulation artifacts were removed by linearly interpolating the signal from 1 ms before to 5 ms after the onset of each stimulus. At this point both ESI and EEG data were band-pass filtered (50–800 Hz, FIR filter). Dorsal electrodes with signal amplitude with a variance >5 times larger than the variance of the mean across channels for at least 50% of the stimulation periods were removed and spline interpolated using the neighbouring electrodes. Across participants, the mean number of interpolated electrodes was 10.3±3.7% of the total number of electrodes.

In the resulting continuous data, somatosensory stimuli that occurred within ±150 ms from the R peak were excluded from the subsequent analysis. These discarded stimuli constituted 33.9±14.9% (range 21.7-45.8%) of the total number of stimuli per subject. The proportion of stimuli discarded depended largely on the participant’s heart rate. The remaining continuous data were thereupon segmented into 300-ms long epochs (-100 to +200 ms) relative to the onset of the stimuli. These epochs were baseline corrected using the -100 to -1 ms interval as a reference, and linearly detrended.

Finally, epochs with peak-to-peak amplitude values exceeding a threshold between 100 and 250 µV (160±44 µV; defined individually according to the signal-to-noise ratio) in at least one electrode (i.e., epochs probably contaminated by artifacts) were rejected. Considering both ESI and EEG datasets, these epochs were 2.2±2.9% of the total number of epochs devoid of ECG artifacts.

The resulting clean epochs were labeled according to the possible combinations of the four experimental conditions (stimulation side: right or left; cognitive task: omission counting, letter memorization).

To investigate the differences in the responses elicited by right and left median nerve stimulation, all epochs obtained during the omission and letter memorization tasks were pooled. This procedure yielded two average waveforms per participant (right and left stimulation). In contrast, to investigate the differences in the responses elicited during the omission vs letter memorization task, all epochs obtained following right median nerve stimulation were flipped with respect to the midline, so that they could be pooled with the data obtained following left median nerve stimulation. This procedure yielded two additional average waveforms per participant (omission and letter memorization). Finally, to investigate the temporal and spatial properties of the response along the somatosensory pathways, the flipped epochs were pooled across tasks, as if the stimuli were always delivered to the left side. This procedure yielded one additional average waveform per participant.

Given that early-latency somatosensory responses are notoriously variable across individuals as a function of arm length and height (Chabot et al., 1985), to minimize inter-individual variability and perform group-level statistics, single-subject average waveforms were peak aligned using a validated procedure based on linear time-warping (Hu et al., 2014; Mancini et al., 2015). Specifically, we aligned (1) eN9 peaks measured at the ipsilateral Erb’s point, (2) sN13 peaks measured at midline dorsal electrode S8, and (3) cN20 and cP25 peaks measured at the contralateral parietal scalp electrode Pc.

After realignment, the single-subject average waveforms were averaged to obtain group-level waveforms. Group-level topographies were computed by spline interpolation.

### Latency maps

To determine whether the recorded components, such as the sP9 and sN13, reflect traveling far-field action potentials or local post-synaptic activity, we computed spatial maps of peak latency distributions (Massimini et al., 2004). We restricted this analysis to the electrodes positioned in the upper-half of the recording array, to exclude electrodes in which the component of interest was absent or extremely small in amplitude. We measured, separately for each component, the peak latency from each electrode, and we subsequently color coded peak latency relative to the electrode with the shortest latency (Figure 4). In these images, traveling action potentials would appear as a smooth and continuous unidirectional gradient. On the contrary, the lack of such a unidirectional gradient would suggest that the component observed in the time-domain reflects local post-synaptic activity.

**Figure 3.**
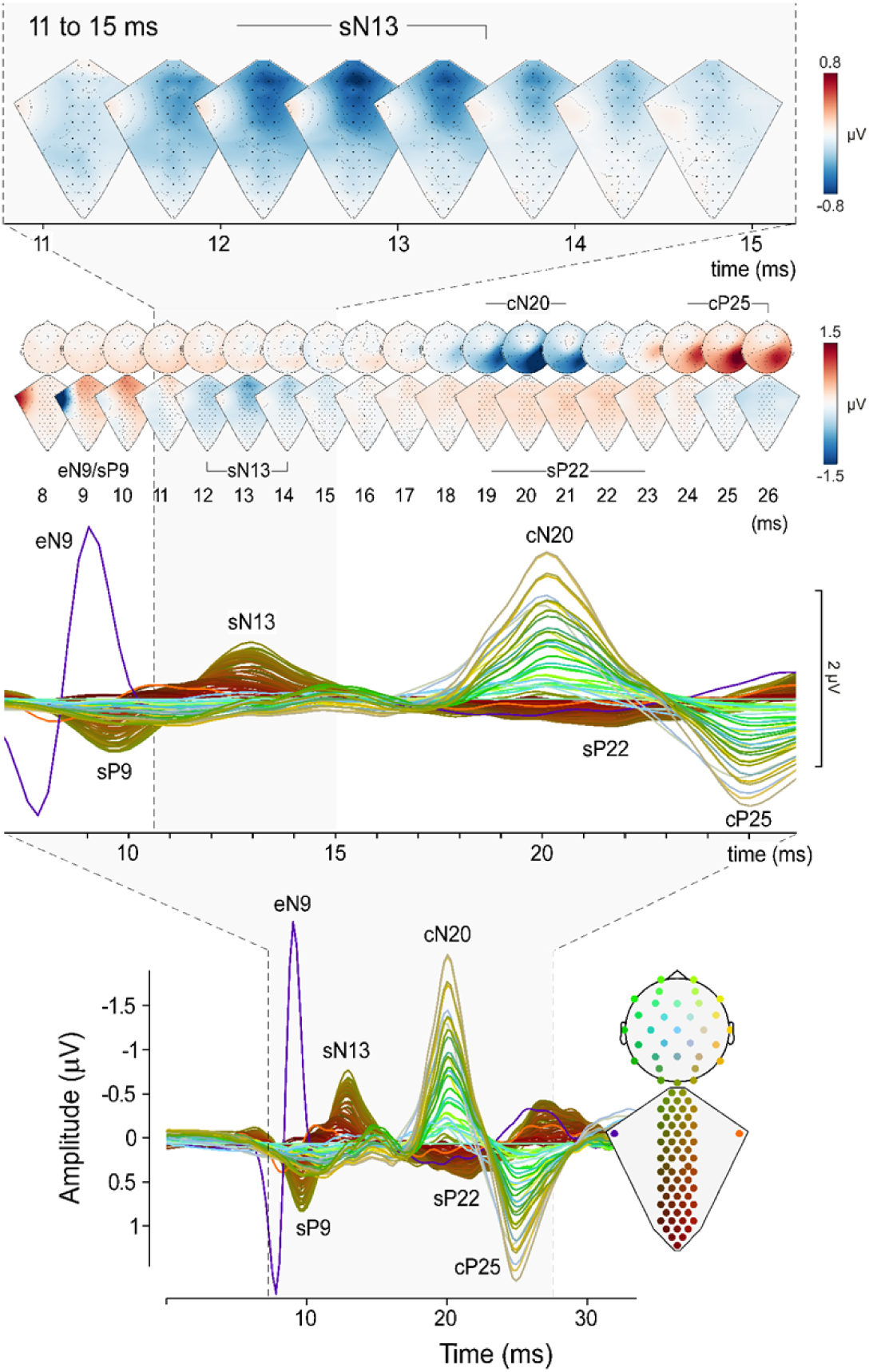
Electral Spinal Imaging (ESI): High-resolution images of spinal cord activity during somatosensory stimulation. *Bottom panel*: Recordings obtained using 97 electrodes (2 electrodes located on the right and left ERB points, 63 dorsal electrodes spanning from the cervical (C2) to the thoracic (T8) segments, and 32 scalp electrodes). The electrode positions shown in the bottom right figurine and their recording of the response elicited by somatosensory stimulation (right) are colour-coded such that neighboring channels have similar colors. Signals were referenced as described in Figure 1. *Middle panel*: Enlargement of the responses between 8 and 26 ms post-stimulus. *Top panel*: Lower row shows dorsal and scalp maps with 1-ms resolution. The eN9, eP9, sN13, p22, cN20 and cP25 peaks are labelled. Upper row enlarges the dorsum maps of spinal cord activity between 11 and 16 ms post-stimulus. Note the peak of post-synaptic activity occurring around 13 ms post-stimulus.

**Figure 4.**
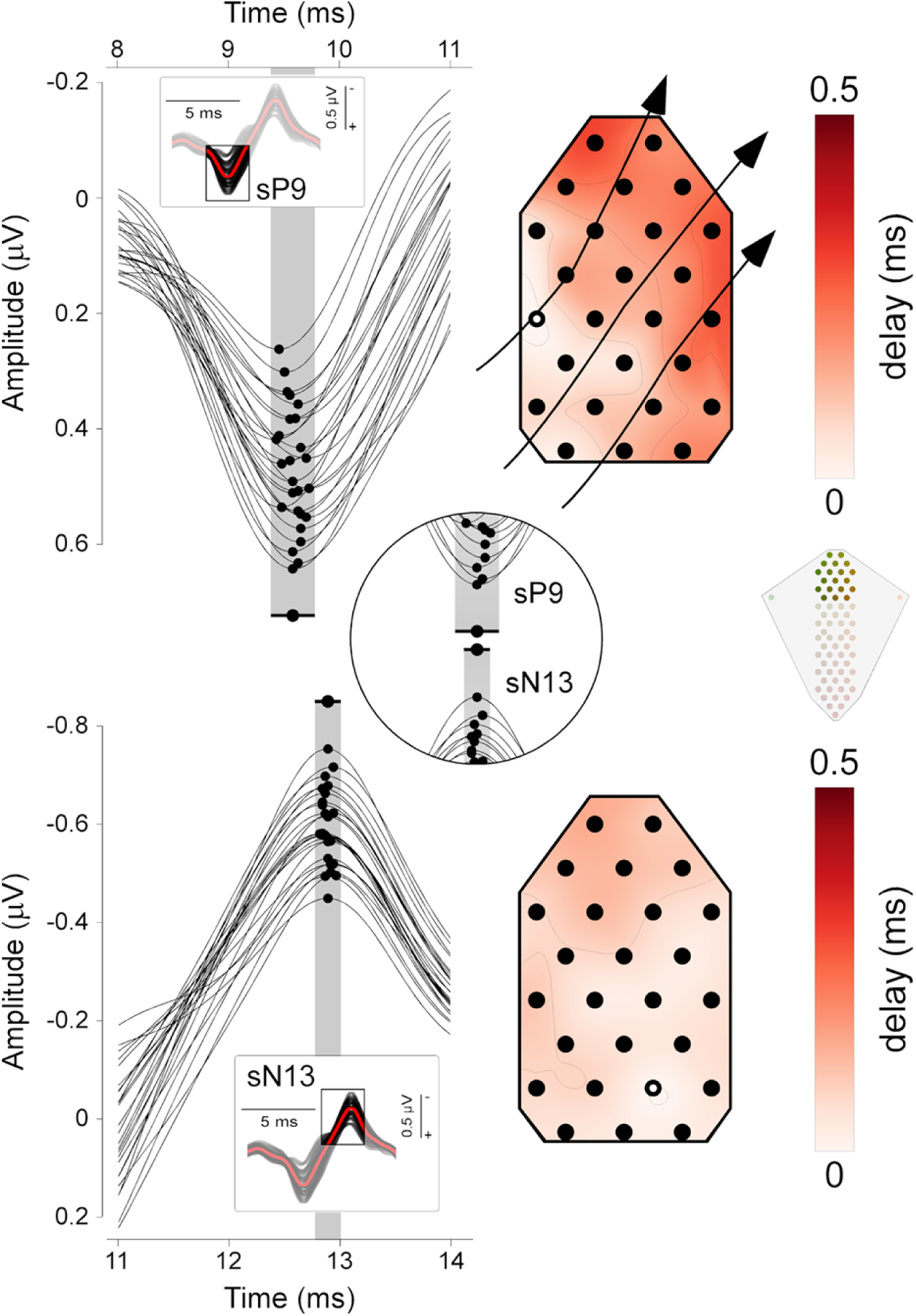
Latency distributions of the sP9 and sN13 waves. Group-level data. *Left column*: sP9 (top) and sN13 (bottom) waves recorded from the top dorsal electrodes. Insets show the response over a larger time window; the box indicates the intervals around sP9 and sN13. Dots indicate the peak latency recorded from each electrode. The red dot and horizontal bar indicate the mean latency and latency range, respectively. *Right column*: Maps of the latency delays, with color-coding of time relative to the electrode with the shortest latency (highlighted in white used as time 0). Note how, for the sP9 wave, the electrodes with shortest (whiter regions of spinal electrode topography, broadly corresponding to the C6-C7 segment) and longest latencies (redder regions) show a clear spatial gradient, indicating that the sP9 wave travels in the lateral-medial and caudal-rostral directions. Compare with the lack of any clear spatial gradient in the sN13.

### Lateralization analysis

To determine whether the EEG and ESI responses were lateralized as a function of the stimulated side, single-subject waveforms elicited by right and left median nerve stimulation were compared using a point-by-point paired sample t-test. The significance level was set at p=0.01. To correct for multiple comparisons, an electrode was considered significant only if (1) it was part of a cluster of at least three adjacent electrodes with a p-value<0.01, and (2) it was part of a time window with an average p<0.01 in at least 4 consecutive time points (i.e. for a duration of at least 1 ms). Group-level difference waveforms (left *minus* right) were calculated to show their spatial distribution. Dorsal and scalp topographies and their differences were plotted at the peak latency of the eN9, sN13, cN20, sP22, and cP25 waves.

### Attentional task analysis

To determine whether the EEG and ESI responses were different as a function of the attentional task, we performed the same procedure described above but comparing the omission counting and letter memorization conditions. Group-level difference waveforms were calculated as letter memorization *minus* omission counting (Figure 6).

## Results

### Response waveforms and topographies

At single-subject and group-level, somatosensory stimuli elicited, clear responses from the brachial plexus (eP8 and eN9), the spinal cord (sP9, sN13, sP22), and the brain (cN20 and cP25) (Figure 1, left panel). By exploiting a high-density spatial sampling, we were able to reveal how the neural activity along the somatosensory pathways unfolds in time and space with sub-millisecond temporal resolution (Figure 2, Video 1).

#### Erb components

We were able to identify the travelling afferent volley at the brachial plexus ipsilateral to the stimulated side, in the form of a positive-negative complex (eP8/eN9). Its average amplitude and duration (calculated, at Erb, spinal, and scalp recordings, as the width at half maximum) were as follows: eP8 2.0±0.9 μV, 1.9±0.5 ms; eN9 -2.9±1.0 μV, 1.8±0.2 ms.

#### Spinal components

Concomitant with the eN9 wave at the brachial plexus, upper dorsal electrodes disclosed a first positive spinal response peaking around 9.5 ms post-stimulus (sP9: 0.93±0.3 μV, 3.2±0.8 ms; channel S8), followed by the sN13 wave (-0.9±0.4 μV, 3.7±0.8 ms; channel S8). Both sP9 and sN13 had a single clear maximum, centered on the dorsal electrodes overlying the spinal segments C4 to C6, with no clear lateralization (Figures 2, 3). After the end of the sN13 wave, there was a period of ∼3 ms with no sign of electrical activity, which was interrupted by the occurrence of a longer-lasting positive wave (sP22: 0.7±0.4 μV, 5.5±1.7 ms; channel S8). sP22 had a maximum over the midline dorsal electrodes overlying the spinal segments C4 to C7. Compared to the earlier spinal responses (sP8 and sN13), sP22 was longer-lasting and more widespread in both the rostrocaudal and the mediolateral directions (Figures 2, 3), suggesting a longer-loop, more polysynaptic generation mechanism compared to the earlier, sharper spinal components.

#### Scalp components

At scalp level, somatosensory stimuli elicited the canonical cN20-cP25 complex, maximal over P3/4 CP5/6 and P7/8, i.e. the electrodes broadly corresponding to the hand representation of the primary somatosensory cortex contralateral to the stimulated side (Nuwer & Dawson, 1984) (Figure 2). The amplitude and duration of the cN20-cP25 complex were as follows: cN20: -2.1±0.9 μV, 5.4±0.7 ms; cP25: 1.6±1.0 μV, 6.5±1.4 ms.

### Temporal dynamics of spinal responses

To gain insight into the nature of the sP9 and sN13 components, and specifically to understand whether they represent far-field traveling action potentials or segmental post-synaptic activity, we built latency delay images, where colors indicate the peak latency delay relative to the electrode with the shortest latency (Figure 4). For the sP9 wave, latency delays followed a clear gradient, whose spatial distribution indicated that the sP9 reflects far-field action potentials entering the spinal cord at ∼C6-C7 level, and travelling in the lateral-medial and caudal-rostral directions (Figure 4, top). Video 2 shows the sP9 sweeping the field of view from the ∼C6-C7 entry level to the most rostral dorsal electrodes. In contrast, the sN13 did not show signs of travelling (Figure 4, bottom).

Taken together, these results suggest that the sP9 mostly reflects the extracellular events associated with the propagation of action potentials of Aß afferent fibers entering the spinal cord from the brachial plexus and the dorsal roots, and then travelling in the dorsal columns. In contrast, sN13 seems to largely reflect post-synaptic activity occurring in the dorsal horn at C4-C6 levels, consequent to the arrival of the Aß afferent input.

### Response lateralization

Figure 5 shows the grand average ESI and EEG waveforms superimposed, together with the dorsal and scalp topographies at the peak latency of the eN9, sN13, cN20, sP22 and cP25 waves, for each stimulation side. At dorsal level, the topography of eN9 elicited by median nerve stimulation displayed a clear negative maximum at the electrode detecting the activity of the brachial plexus ipsilateral to the stimulated side. We observed a larger eN9 amplitude in the ERB_ipsi_ compared to the ERB_contra_ electrode (p=0.0008, paired-sample t-test). In contrast, the topography of the sN13 and sP22 waves was similar following right and left stimulation, with no electrodes showing significant differences (p-value range = 0.066 – 0.906).

**Figure 5.**
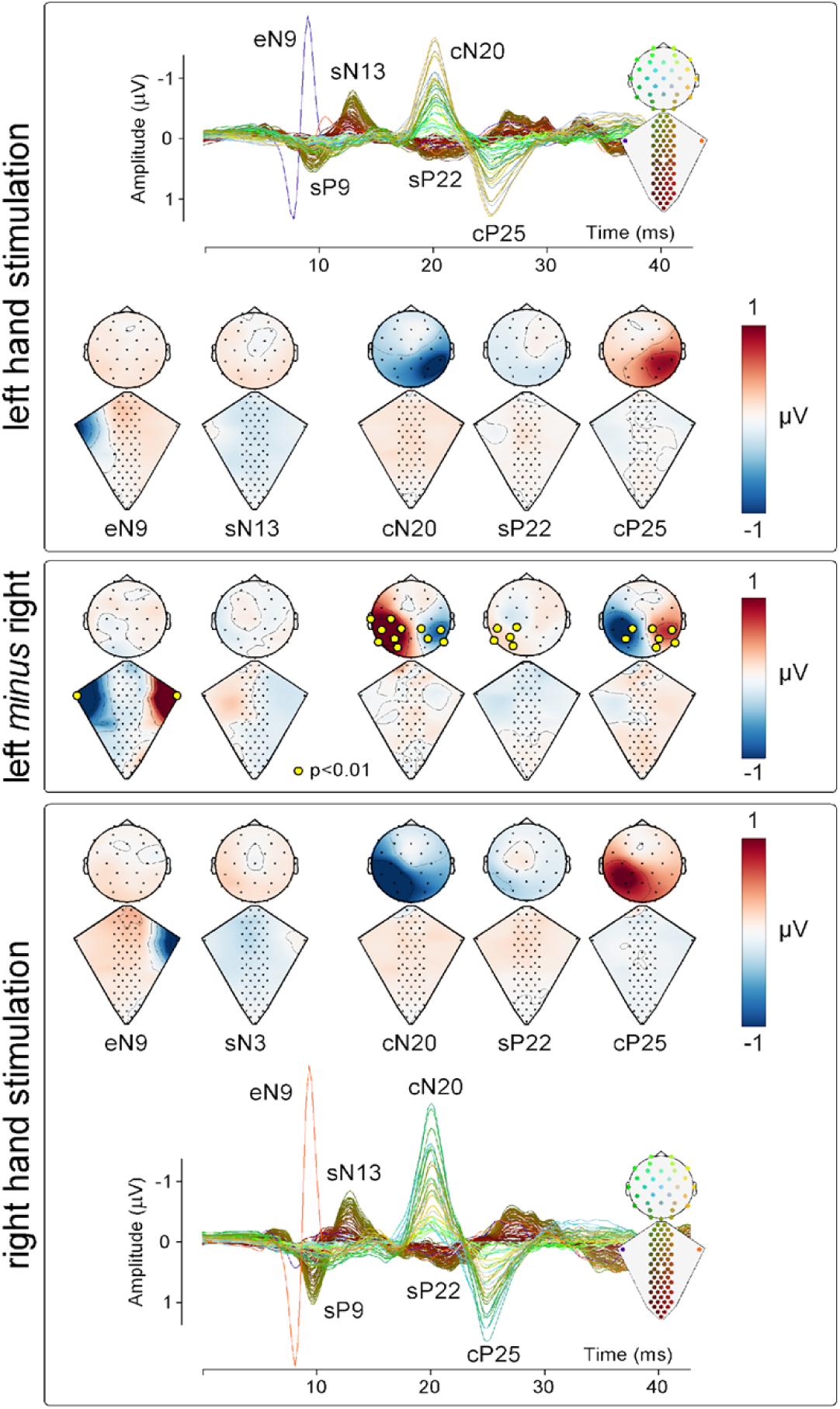
Lateralization of neural responses along the somatosensory pathways. *Top and bottom panels*: Neural responses recorded along the somatosensory pathways following the stimulation of the median nerve at the left and right wrist, respectively. Top and bottom panels display the group-level response superimposed from all 97 electrodes superimposed, referenced to S64 for the dorsal electrodes, and to Fz for the scalp electrodes. Electrode position and responses are colour-coded according to the scheme shown in the inset. These panels also contain dorsum and scalp maps at the latency of the eN9, sN13, cN20, sP22 and cP25 peaks. *Middle panel*: Difference maps obtained by subtracting the right response from the left response, at the same latencies as the maps shown in the top and bottom panels. Note the clear lateralization of the response recorded from the brachial plexus (eN9) and the brain (cN20-cP25), and the lack of lateralization of the spinal responses (sN13 and sP22).

At scalp level, the topographies of both the cN20 and cP25 waves were clearly lateralized. The cluster of electrodes overlying the primary somatosensory cortex contralateral to the stimulated side displayed higher amplitudes compared to the ipsilateral side (cN20: mean p-value=0.004, p-value range=0.0002–0.0096; cP25: mean p-value=0.0011, p-value range=1.5e-7 – 0.005; paired-sample t-test).

### Attentional modulation

Figure 6 shows, for each attentional condition, the grand average ESI and EEG waveforms superimposed, together with the dorsal and scalp topographies at the peak latency of the eN9, sN13, cN20, sP22 and cP25 waves. We found strong evidence for a larger sP22 amplitude in the letter memorization condition (mean p-value=0.003, p-value range=0.0001-0.0099; paired-sample t-test). In contrast, the eN9 and sN13 amplitudes were similar during the two attentional tasks, with not enough electrodes showing significant differences (p-value range=0.061–0.896). Similarly, there was no evidence of any difference in the amplitude of the cN20 and cP25 (cN20: p-value range=0.0463–0.722; cP25: p-value range=0.060–0.856; paired-sample t-test).

**Figure 6.**
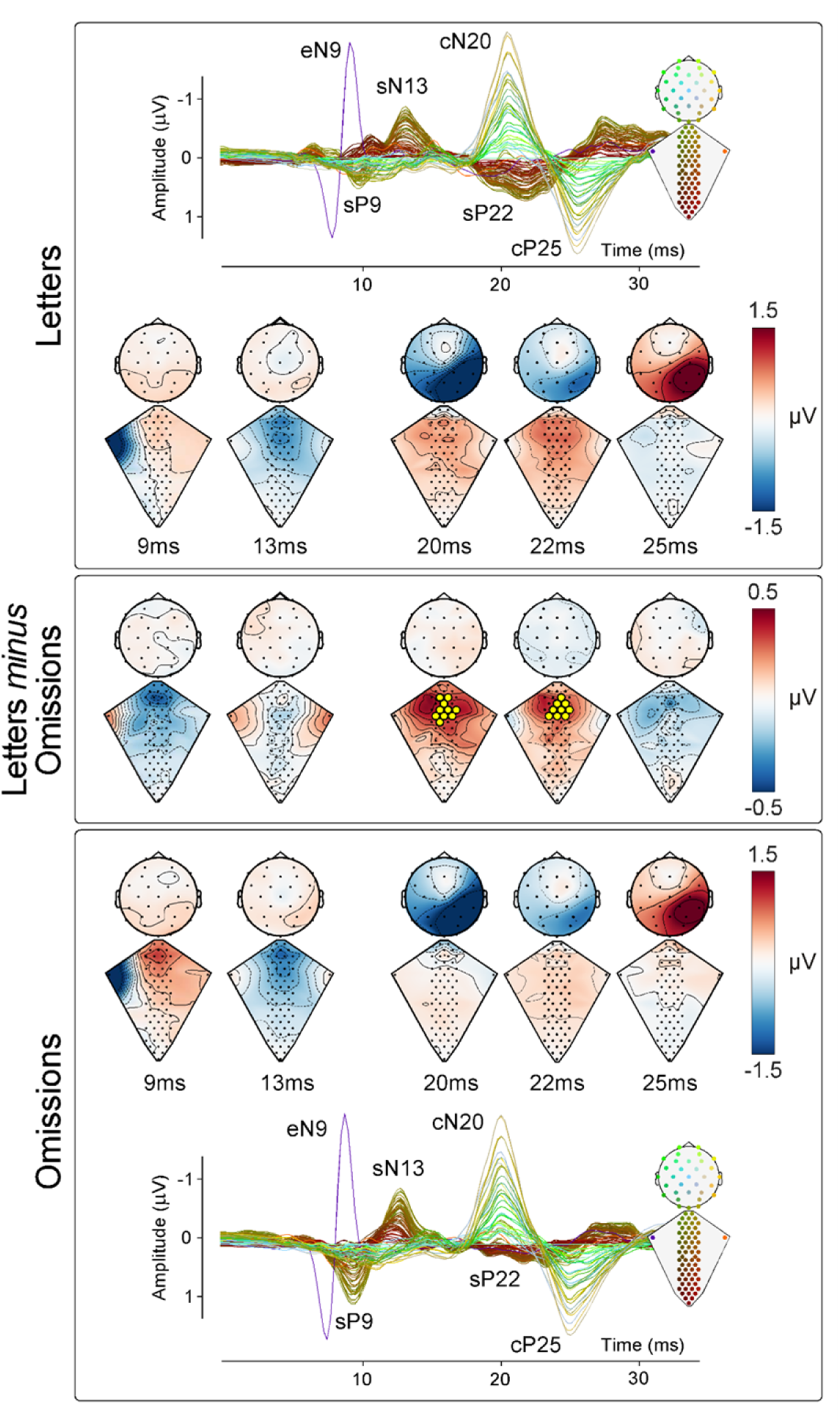
Attentional modulation of spinal cord activity. *Top and bottom panel*: Neural responses recorded along the somatosensory pathways under the conditions of letter memorization (top panel), and attention to somatosensory stimulation (counting omissions, bottom panel). Both panels show the group-level response distribution on the dorsum and scalp at the eN9, sN13, cN20, and cP25peak latencies. Electrode position and responses are colour-coded according to the scheme shown in the inset. *Middle panel*: Difference maps obtained by subtracting the responses elicited during the omission task from those elicited during the letter memorization task, at the same latencies as the maps in the top and bottom panels. Electrodes where this difference was significant are highlighted in yellow.

## Discussion

In this work, we describe a novel recording and analysis approach to obtain images of the electrical activity of the spinal cord non-invasively and with high temporal and spatial resolution (Electro-Spinal Imaging, ESI). Crucially, this approach does not entail any form of template-matching strategies (such as canonical correlation analysis, CCA; Li et al., 2009; Nierula et al., 2024; Woody, 1967), which are prone to generate the false belief of consistent responses (Mayhew et al., 2006).

Exploiting this electro-spinal imaging approach (ESI), we provide a detailed description and physiological characterization of the spatiotemporal dynamics of the peripheral, spinal and cortical activity elicited by somatosensory stimulation. Importantly, we demonstrate how cognition can modulate post-synaptic activity at spinal cord level.

We obtained four key results regarding spinal cord activity (1) We identified three distinct responses: sP9, sN13 and sP22. (2) The sP9 is a traveling wave, while the sN13 and sP22 reflect local, post-synaptic activity. (3) While the sP9 response is first seen on the dorsal electrodes ipsilateral to the stimulated side, the sN13 and sP22 were not clearly lateralised with respect to the side of stimulation. (4) Attention strongly modulates the amplitude of the sP22, but not that of the sP9 and sN13.

### Bipolar shape of the eP8/eN9 complex: comparison with the tripolar shape of near nerve recordings

With the electrodes located at the Erb point we recorded the classical travelling afferent volley at the brachial plexus ipsilateral to the stimulated side as a positive-negative bipolar eP8/eN9 complex (Figure 2, 3, Video 1). This pattern contrasts with the classical descriptions of a triphasic positive-negative-positive pattern recorded when action potentials travel along nerve fibers (e.g.: Buchthal & Rosenfalck, 1966; Lorente de No, 1947) This difference may arise because classical monopolar extracellular nerve recordings use an active electrode detecting the depolarisation occurring along the fibre: the active electrode sees first the approaching sink (first positivity), then the sink passes under the electrode (large negative deflection) and finally the sink goes away (second positivity) (Lorente de No, 1947). In contrast, our recordings showed a biphasic, not a triphasic pattern. This biphasic pattern may reflect the so-called *end potential* phenomenon, in which the second positive component is attenuated because the “going away” phase escapes the detection field of the electrode, due to either a change of direction of the axon or to an end synapse (Jewett & Deupree, 1989).

We now discuss each of the three components that we identified in dorsal recordings (sP9, sN13, and sP22; Figures 1, 2; Video 1).

### The sP9 reflects somatosensory input entering the cord through the dorsal roots

Invasive and non-invasive studies in a number of species, including rodents, human and non-human primates also described that the earliest spinal response following somatosensory stimulation is a positive deflection (labelled as P1 or P9) (Dimitrijevic & Halter, 1995). Invasive experiments using serial recordings at different locations along the cord convincingly identified the origin of this component in the afferent volley travelling through the dorsal roots into the spinal cord (Campbell, 1945). The high-spatial resolution of ESI allowed us to investigate this travelling property of the sP9 component directly. Our latency delay images confirmed that the sP9 wave first appears in the lateral electrodes ipsilateral to the stimulated side at C6-C7 level, and then travels along both lateral-medial and caudal-rostral directions (Figure 4: Video 1). Thus, the sP9 likely reflects the current sink entering the cord through the dorsal roots and travelling rostrally along the dorsal column tracts (Figure 7).

**Figure 7.**
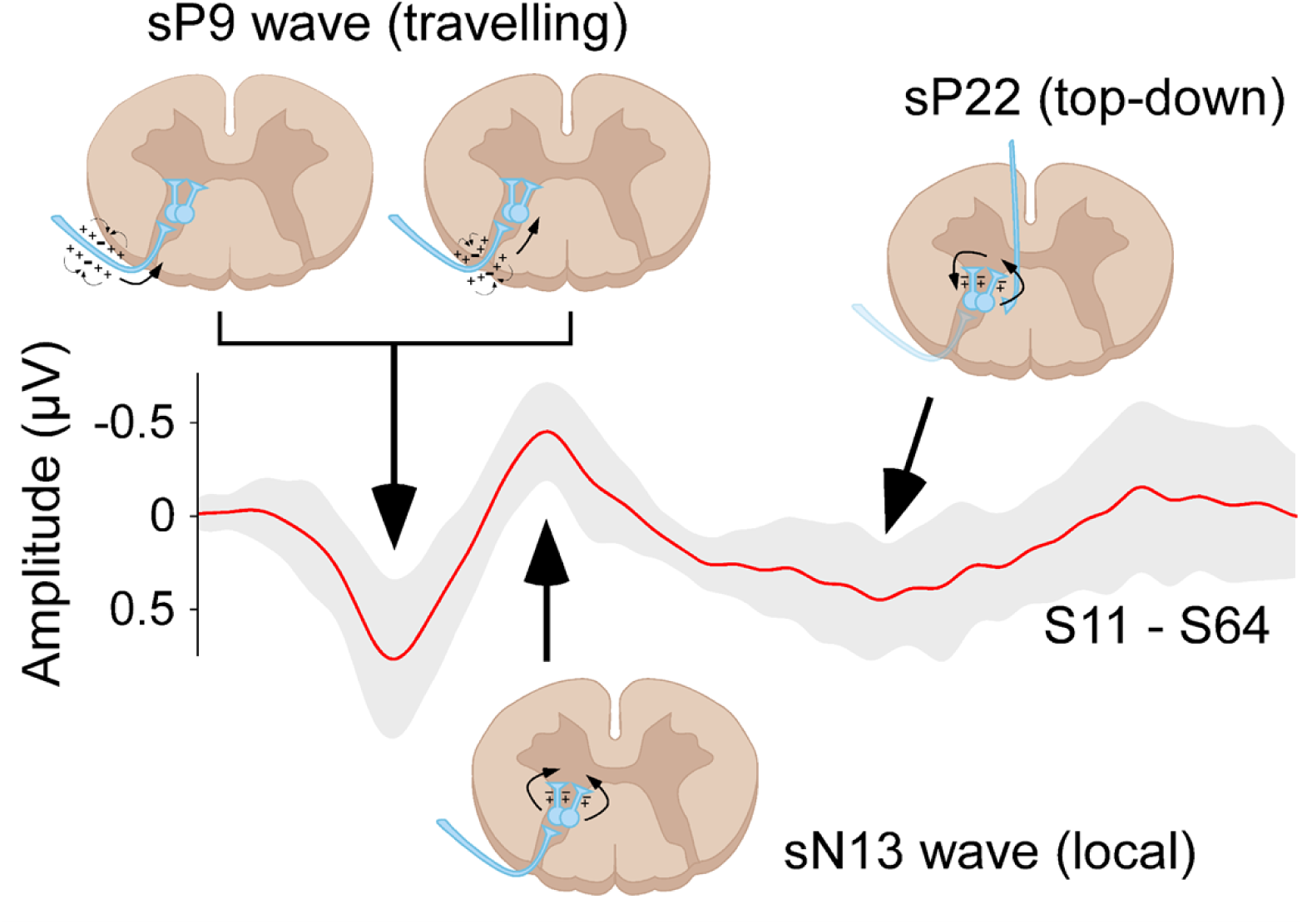
Neural generators of the recorded responses. Schematic representations of the putative spinal mechanisms generating the electrocortical responses recorded with ESI. The sP9 appears as a travelling wave (see also the delay map in Figure 4) as it reflects the current sink entering the cord through the dorsal roots and travelling rostrally along the dorsal column tracts. In contrast, the stationary sN13 reflects the segmental post-synaptic activity occurring in the deep dorsal horn following the first arrival of somatosensory input. Finally, the late sP22 is not directly consequent to a local, segmental effect of the incoming somatosensory input, but instead reflects stimulus-triggered activation of a long-loop circuit involving supraspinal structures that, in turn, project top-down to the spinal cord. For this reason the sP22 encompasses spinal segments rostral and caudal to those where the sP9 and sP13 are recorded (Figure 3, Video 1).

### The sN13 reflects segmental post-synaptic activity in the dorsal horn

The sP9 was followed by the sN13 (Figures 2, 3). In striking contrast with the sP9, the sN13 was stationary: while also appearing on the dorsal electrodes overlying the C6-C7 segments, it lacked the traveling-wave profile that characterized the sP9 (Figure 4, Video 1). Thus, the sN13 likely reflects post-synaptic activity occurring locally, at the level of the dorsal horn. Our sN13 is thus equivalent to the so-called N1 recorded both subdurally and intraparenchymally in cats, dogs, humans, and non-human primates (Beall et al., 1977; Bernhard & Widén, 1953a; Coombs et al., 1956; De Molina & Gray, 1957; Eccles & Sherrington, 1930; Gasser & Graham, 1933). The observations that the N1 is measured (1) already in response to low stimulus intensities activating large-diameter A-afferents (Beall et al., 1977), (2) maximally at the segmental level that corresponds to the stimulated dermatome (Willis et al., 1973), and (3) at the same segmental level as the traveling sP9, all indicate that the sN13 reflects the post-synaptic activity occurring in laminae IV and V. Thus, we take the sN13 observed in our recordings as a non-invasive readout of the post-synaptic activity occurring in the deep dorsal horn following the first arrival of somatosensory input (Figure 7). Given the sP9-sN13 latency and the average length between stimulation site and recording electrodes of ∼70 cm, this input travels along medium-diameter type-II Aβ afferents conducting at approximately 50-70 m/s.

### The sP22 reflects top-down supraspinal projections

The latest spinal component sP22 was also maximal on the electrodes overlying C4-C7 spinal segments. Compared to the preceding sP9 and sN13, it was longer-lasting and more widespread in both rostrocaudal and mediolateral directions (Figure 2, 3).

In animal studies the P1-N1 waves (corresponding to our sP9-sN13) are immediately followed by a lower-frequency positivity that is typically divided into two distinct components: a ‘fast’ P2f and a ‘slow’ P2s. The P2f, which is only observed in invasive recordings (Jeanmonod et al., 1991) and resists spinal transection (Bernhard & Widén, 1953b), is interpreted as reflecting a primary afferents depolarization (Carpenter et al., 1963). The subsequent P2s, in contrast, depends on the animal’s behavioural state and is abolished by spinal transection, indicating that it reflects the consequences of top-down supraspinal projections (Shimoji & Kano, 1975; Tang, 1969).

Several pieces of evidence indicate that our sP22 corresponds to the top-down driven P2s of the animal invasive literature: first, its relatively long latency is compatible with the top-down hypothesis arising from animal experiments (Figure 2). Second, the sP22 is the only spinal component that was clearly modulated by our cognitive task (Figure 6). Interestingly, the spatial resolution offered by high-density ESI, allowed us to observe that the sP22 encompasses spinal segments both rostral and caudal to those where the sP9 and sP13 are located (Figure 3). This suggests that sP22 is not a direct consequence of the incoming somatosensory signals – since these would be expected to become evident more rostrally than caudally. Instead, sP22 may reflect stimulus-triggered activation of top-down supraspinal projections (Figure 7). Indeed, only afferents belonging to small-myelinated Aδ and unmyelinated C units travel both rostrally and caudally in the Lissauer’s tract after entering the dorsal root, and thus elicit responses (Porro & Cavazzuti, 1992) and perceptual effects (Mitchell et al., 2024) not strictly localized at the level of the dermatomal stimulation. Crucially, these afferents are not activated by the electrical stimulation we used, which preferentially activates large-myelinated, fast-conducting Aβ fibres that project rostrally rather than caudally. Further, the afferent volley from slower fibres does not reach the spinal cord until 35-45 ms after the electrical stimulation. Thus, the combination of the temporal (22 ms latency) and spatial (similar rostral and caudal distribution) properties of the sP22, suggesting a polysynaptic generation mechanism, points towards a supraspinal rather than an afferent origin.

### Response lateralization

Consistent with findings reported in the literature, we observed significant evidence of response lateralization at both the Erb’s point and the scalp (Nuwer & Packwood, 2008). At the brachial plexus, the eP8 and eN9 responses were detected only on the side ipsilateral to the stimulation, whereas at the scalp, the cN20 and cP25 responses appeared exclusively contralateral to the stimulation (Figure 5).

Previous studies using both invasive recordings in non-human primates and intraoperative monitoring of SEPs in humans have reported the possibility of lateralized spinal responses following median nerve stimulation. More specifically, in anesthetized cynomolgus monkeys the origin of the negative components comparable to the human sN13 is in the laminae IV-VI of the dorsal horn ipsilateral to the stimulation side (Beall et al., 1977). We therefore took advantage of the topographical information provided by ESI to test whether the responses we recorded were similarly lateralized. To the best of our knowledge, we are the first to perform a point-by-point comparison between right and left stimulation across the full time-course of the stimulus-evoked response. The advantage of performing paired t-tests along the whole time course allows for the identification of effects outside of the temporal regions of interest, such as before or after the peaks of the components. We found no statistical evidence of lateralization at any point during the evoked response (Figure 5). We cannot exclude the possibility that the neural response may in fact be lateralized, but that factors such as low spatial separation of electrodes along the mediolateral axis, effects of volume conduction, and low signal-to-noise ratio made our recordings unable to show reliable evidence for lateralization.

### Attentional modulation

Cortical top-down control appears to be a general principle of neural organization (Colloca & Barsky, 2020; Fields, 2004). Such processes may strongly affect neural activity in subcortical structures, including the spinal cord (e.g. Seki & Fetz, 2012).

For example, functional MRI studies of the spinal cord have shown that nocebo manipulations enhance pain-related BOLD activity in the dorsal horn ipsilateral to a nociceptive stimulus, This suggests a tonic top-down pain-facilitation mechanism occurring at spinal cord level (Geuter & Büchel, 2013). Similarly, nociceptive stimuli delivered during high-load cognitive tasks result in lower pain ratings and weaker BOLD responses in the ipsilateral dorsal horn, compared to a low-load control condition (Sprenger et al., 2012). However, given that BOLD activity integrates neural activity across several seconds (Logothetis et al., 2001), fMRI cannot unravel the temporal dynamics of these modulations. By combining an attentional paradigm with the high temporal resolution of ESI, we were able to show that attention selectively influences specific components of the spinal response (Figure 6). Specifically, we found that the sP22 amplitude (which, as discussed above, is the human equivalent of the P2s recorded in animals and likely reflects stimulus-triggered activation of top-down supraspinal projections) depends strongly on current cognitive task and focus of attention, while earlier spinal and cortical responses do not. These results are informative about the underlying nature of the attentional modulation that can only be coarsely measured with spinal fMRI. In particular, our results suggest that attention does not cause a *general* gating of all somatosensory-evoked activity at spinal cord level. Instead, attentional effects may be specifically linked to supraspinal modulation of synaptic activity within the spinal cord.

---

**Video 1.** ESI experimental setup and responses: [Version A] [Version B]

**Video 2.** sP9 wave travelling [Video 2]

